# Sound level context modulates neural activity in the human brainstem

**DOI:** 10.1101/2021.06.24.449846

**Authors:** Björn Herrmann, Sonia Yasmin, Kurdo Araz, David W. Purcell, Ingrid S. Johnsrude

## Abstract

Optimal perception requires adaptation to sounds in the environment. Adaptation involves representing the acoustic stimulation history in neural response patterns, for example, by altering response magnitude or latency as sound-level context change. Neurons in the auditory brainstem of rodents are sensitive to acoustic stimulation history and sound-level context (often referred to as sensitivity to stimulus statistics), but the degree to which the human brainstem exhibits such neural adaptation is unclear. In six electroencephalography experiments with over 125 participants, we demonstrate that the response latency of the human brainstem is sensitive to the history of acoustic stimulation over a few tens of milliseconds. We further show that human brainstem responses adapt to sound-level context in, at least, the last 44 ms, but that neural sensitivity to sound-level context decreases when the time window over which acoustic stimuli need to be integrated becomes wider. Our study thus provides evidence of adaptation to sound-level context in the human brainstem and of the timescale over which sound-level information affects neural responses to sound. The research delivers an important link to studies on neural adaptation in non-human animals.

## Introduction

Optimal sound perception requires an individual to adapt flexibly to their changing acoustic environments^1–5^. One crucial mechanism underlying such flexibility is neural adaptation. Neural adaptation describes a decrease in neural response magnitude and/or an increase in response latency with sustained or repeated acoustic stimulation^6–8^. Adaptation provides a means by which the history of acoustic information is represented in neural circuits, which, in turn, influences neural responses to future sounds. Through adaptation, neural circuits represent when a sound has occurred in time^9–11^ as well as statistical properties of entire acoustic feature dimensions, such as spectral variance^12–14^ or the modal/mean sound level of acoustic environments^15–17^.

Work in rodents demonstrates that auditory nerve fibers and neurons in the brainstem and auditory cortex are capable of adapting to ambient sound-level contexts, often referred to as adaptation to sound-level statistics^15,18–23^. Work in humans has documented adaptation to sound-level context and sensitivity to violations of expected sound levels in primary and secondary auditory cortices^16,24–27^. However, it is currently unclear whether neurons at lower stages of the human auditory system, such as the brainstem, adapt to ambient sound-level information and, if so, over what timescale sound-level information is represented in neural responses. In a series of electroencephalography (EEG) experiments, we evaluate neural adaptation in the human brainstem to contextual information of acoustic environments.

Investigations of neural adaptation often rely on manipulations of the interval duration between two sounds^10,28^. The longer the interval between two sounds, the more time neurons have to recover from adaptation after responding to the first sound. Accordingly, responses to the second sound are faster and larger as the interval between the two sounds increases^9–11,29–34^. Adaptation of human brainstem responses appears to manifest most prominently in the response latency, while the response amplitude appears to adapt less^35–38^; but see^33^.

When neural response properties change in response to recent stimulation, this may reflect simply the interval since the last sound, or it may reflect something more complex about the history of stimulation, spanning a period incorporating several sounds. The latter case is more relevant for adaptation to sound-level context. Some previous studies have demonstrated that the latency of human brainstem responses adapts to the stimulation history beyond the influence of the directly preceding sound^28,39,40^. In a few instances, human brainstem activity has been shown to integrate previous sound stimulation over several hundreds of milliseconds^28,39^, suggesting that the human brainstem may be capable of representing sound-level information.

The current study investigates whether brainstem responses in humans, similar to rodents^15,21,22^, adapt to sound-level context. In order to investigate neural adaptation to sound-level context one must first establish the extent to which acoustic stimulation history is represented in brainstem responses, given a specific experimental setup. For example, human brainstem neurons demonstrate adaptation over a longer windows for high-compared to low-level sounds^35,41^. Accordingly, we first conduct three EEG experiments (two detailed in the Supplemental Materials) to establish with our experimental setup to what extent brainstem responses in human listeners adapt to stimulation history. We use regular, predictable sound sequences to investigate adaptation of brainstem responses (two experiments), and then use irregular, unpredictable sound sequences (one experiment) to exclude the possibility that prediction-related processes (and not neural adaptation) underpin sensitivity of brainstem responses to acoustic stimulation history. These three experiments enable us, in three subsequent experiments, to investigate whether neurons in the human brainstem adapt to sound-level context. This will reveal the timescale over which sound-level information is represented in human brainstem responses.

## Results

### Human brainstem activity is sensitive to stimulation history incorporating several sounds

A prerequisite for adaptation to sound-level context is the capacity of neurons to represent the sound history over several stimuli in their activity patterns. To investigate whether the human brainstem is sensitive to sound history, we adapted a paradigm that has been successfully utilized to investigate adaptation of the human auditory cortex^42,43^. In Experiment I, normal-hearing younger adults (N=17) listened to click sequences presented at 60 dB sensation level (SL) that continuously “accelerated” and “decelerated”: the click onset-to-onset interval decreased logarithmically from 60 ms to 10 ms and then increased back to 60 ms (Figure 1A). Analyses focused on the Wave V potential at a latency of around 6.6 ms (Figure 1B), which is known to originate from the brainstem^41,44–47^. Wave V amplitude and latency were binned by the preceding onset-to-onset interval, and responses were averaged within bins. Wave V amplitude increased (t_16_ = 2.421, p = 0.028, r_e_ = 0.518) and latency decreased (t_16_ = −9.621, p = 4.7×10^-8^, r_e_ = 0.923) with increasing time since the previous click, as expected (Figure 1B/C).

**Figure 1:**
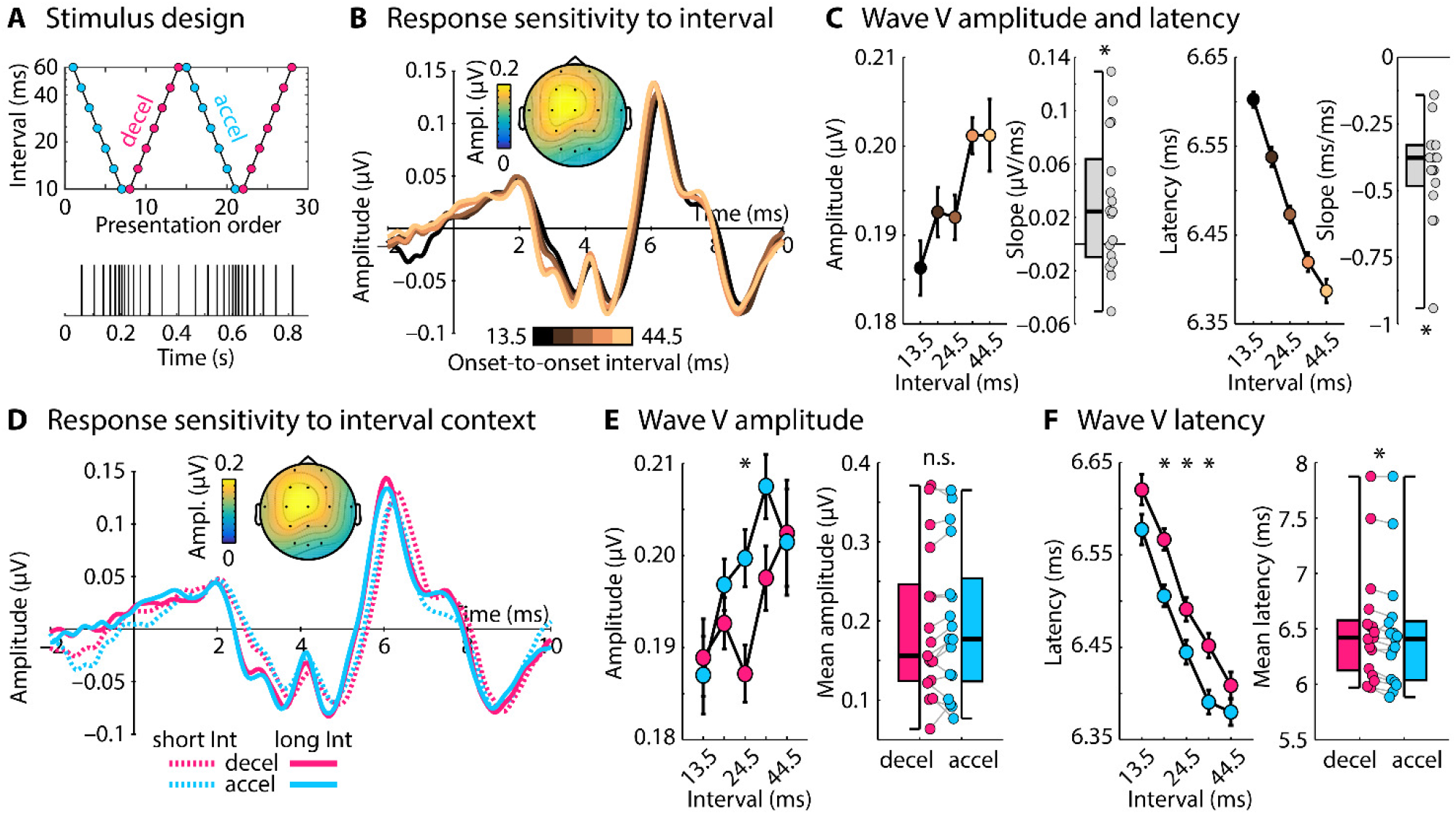
Stimulus design and results for Experiment I. **A:** Onset-to-onset intervals (logarithmic y-axis) of accelerating and decelerating click sequences. **B:** Response time courses to clicks, separately for different onset-to-onset intervals to a preceding click. Topographical distribution reflects the mean Wave V (^~^6.6 ms). **C:** Wave V amplitude and latency as a function of onset-to-onset interval. Box plots and individual data points (dots) reflect the slope from a linear function fit relating Wave V amplitude/latency to onset-to-onset intervals. Wave V latency and amplitude were sensitive to the inter-click-interval (slope significantly different from zero, p ≤ 0.05). **D:** Response time courses for a short (13.5 ms) and a long interval (44.5 ms), separately for decelerating and accelerating sequence contexts. Topographical distribution reflects the mean Wave V. **E/F:** Wave V amplitude and latency as a function of onset-to-onset interval, separately for decelerating and accelerating contexts. Box plots and individual data points (dots) reflect the mean difference between amplitudes/latencies for decelerating and accelerating contexts. Amplitudes were larger and latencies smaller in accelerating compared to decelerating contexts (p ≤ 0.05), demonstrating longer-term influences of the acoustic stimulation on the Wave V response. Error bars reflect the standard error of the mean (removal of between-subject variance^48^). *p ≤ 0.05, n.s. – not significant.

In order to investigate the influence of stimulation history – beyond the interval directly preceding a click – on brainstem responses, trials were separated into accelerating and decelerating sequence contexts (Figure 1D). Both accelerating and decelerating contexts comprise the same onset-to-onset intervals and, as a result, any differences in brainstem responses between contexts could only arise from the longer-term stimulation history: clicks in the accelerating context are preceded by fewer clicks in the same time window compared to the decelerating context, despite the same interval directly preceding any given click. Across stimuli with different preceding intervals, Wave V latency, but not amplitude, was smaller in the accelerating compared to the decelerating context (latency: F_1,16_ = 17.831, p = 6.5×10^-4^, n_p_^2^ = 0.527; amplitude: F_1,16_ = 1.23, p = 0.284, n_p_^2^ = 0.071; Figure 1E/F), demonstrating the influence of longer-term stimulation history on brainstem activity. No interaction between onset-to-onset interval and context (accelerating, decelerating) was observed for either latency or amplitude (ps > 0.05), suggesting that the stimulation history of at least several tens of milliseconds is represented in human brainstem responses (Figure 1F).

We replicated the effects of stimulation history on Wave V latency in Supplementary Experiment A (N=28; using data from a previous study with a different focus^49^) using accelerating-decelerating click sequences with onset-to-onset intervals ranging between 4 ms and 40 ms (Figure S1; Supplementary Materials).

The effect of context on response latency may be due to the density of clicks in an extended time period preceding the click for which the response was analyzed, or it may be related to some kind of prediction, given the orderly presentation of stimuli we used. In Supplementary Experiment B (N=19; Supplementary Materials), we pseudo-randomly vary onset-to-onset intervals between 4 ms and 40 ms to examine whether the Wave V latency is indeed affected by click density in a given time period (Figure S2). Click density had the same effect in this experiment as in the accelerating/decelerating experiment, suggesting that Wave V latency changes are not due to any prediction-related process.

In sum, the results of Experiment I and Supplementary Experiments A and B show that the latency of brainstem activity (Wave V) in response to a click is strongly influenced not only by the interval directly preceding a click, but also by click density before that interval (see also^28,39^). In our data, preceding sound information appears to be integrated over at least several tens of milliseconds (Figure 1F). Our data are consistent with neural adaptation underlying changes in brainstem response latency. Such influence of sound history on neural responses should enable the brainstem to be sensitive to sound-level context.

### Human brainstem activity is affected by sound-level context

In Experiment II, we investigated whether neural activity in the human brainstem is sensitive to sound-level context. To this end, we adapted a paradigm that we have recently used to investigate adaptation to sound-level context in the human auditory cortex^16,24^. Young, normal hearing adults (N=18) listened to clicks presented every 9 ms in two contexts (drawn from two statistical distributions) that differed in the modal sound level: In one context, the modal level was 25 dB SL, whereas in the other context, the modal level was 55 dB SL (Figure 2A). Critically, the sound level of a small set of target clicks (ranging from 15 to 65 dB SL) were identical in both contexts (black dots in Figure 2B/C) as was the level of the click directly preceding each target click (i.e., N-1). Because target and N-1 preceding clicks were identical in the 25-dB and the 55-dB contexts, any differences in neural responses must be due to the longer-term sound-level context of the stimulation. Previous paradigms utilized to investigate adaptation to sound-level context – referred to as adaptation to sound-level statistics – have allowed for the local history (i.e., N-1 context) to contribute to response differences between contexts^15,18–21^. In the current study, in contrast, response differences between sound-level contexts must be due stimulation history beyond the immediate history^16^.

**Figure 2:**
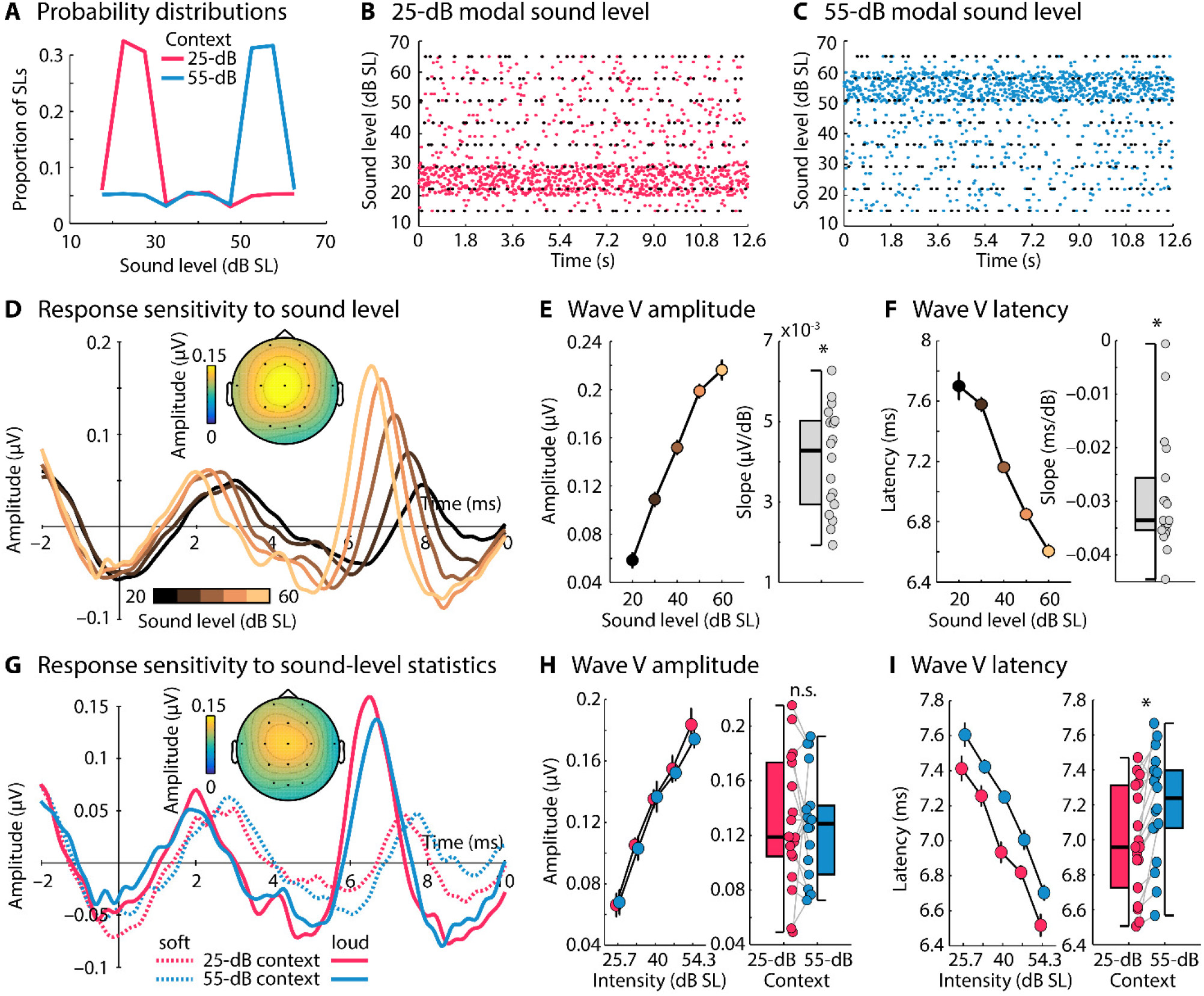
Stimulus design and results for Experiment II. **A:** Example of the two sound-level contexts (distributions with either a 25-dB or a 55-dB modal sound level) utilized to investigate adaptation to sound-level context. **B/C:** Examples of sound-level stimulation in the two contexts. Each dot represents the sound level of one click. The black dots represent the sound level of clicks that were identical in both contexts and for which the sound level of the directly preceding click was also identical in both contexts. **D:** Response time courses to clicks were binned according to their sound level. The topographical distribution reflects the mean Wave V. **E/F:** Wave V amplitude and latency as a function sound level (binned into five categories). Box plots and individual data points (dots) reflect the slope from a linear function fit relating Wave V amplitude/latency to sound level. **G:** Response time courses for low (25.7 dB SL) and high sound levels (54.3 dB SL), separately for the sound-level context with a mode of 25 dB SL and a mode of 55 dB SL. The topographical distribution reflects the mean Wave V. **H/I:** Wave V amplitude and latency as a function sound level, separately for the 25-dB and the 55-dB context. Box plots and individual data points (dots) reflect the mean difference between amplitudes/latencies for the 25-dB and the 55-dB context. Latencies were smaller in the 25-dB compared to the 55-dB context (p ≤ 0.05), demonstrating adaptation to sound-level context in the human brainstem. Error bars reflect the standard error of the mean (removal of between-subject variance^48^). *p ≤ 0.05, n.s. – not significant.

We first confirmed that the sound level of clicks modulates the Wave V. All trials, independent of sound-level context, were binned into five sound-level categories (centered on 20, 30, 40, 50, and 60 dB SL) and averaged. Figure 2D depicts the time courses for each sound-level category. A linear function was fit to Wave V amplitude and latency as function of sound level, separately for each participant, and the resulting linear coefficient was tested against zero. Consistent with previous work, Wave V amplitude increased (t_17_ = 13.289, p = 2.1×10^-10^, r_e_ = 0.955; Figure 2E) and Wave V latency decreased (t_17_ = −11.061, p = 3.5×10^-9^, r_e_ = 0.937; Figure 2F) with increasing sound level^32,41,50,51^.

Next, we limited the analysis to target clicks – that is, the clicks for which level, and the level of the immediately preceding click were identical in both sound-level contexts (25-dB and 55-dB contexts; black dots in Figure 2B). A repeated-measures ANOVA (rmANOVA) revealed that the Wave V amplitude increased with click level, consistent with the previous analysis (F_4,68_ = 59.103, p < 0.001, n_p_^2^ = 0.777), but there was no effect of context nor an interaction between target click level and context (both ps > 0.5; Figure 2H). Wave V latencies were earlier for clicks presented at a high compared to a low level, consistent with the previous analysis (F_4,68_ = 76.544, p < 0.001, n_p_^2^ = 0.818) and were also earlier for clicks presented in the 25-dB compared to the 55-dB context (F_1,17_ = 18.077, p = 5.4×10^-4^, n_p_^2^ = 0.515). Again, no interaction was observed between target click level and context (p > 0.5; Figure 2I). Experiment II shows that the latency, but not amplitude, of the human brainstem Wave V is sensitive to sound-level context.

### Sensitivity of the brainstem to sound-level context decreases with longer intervals between stimuli

In Experiment II, we used an onset-to-onset interval of 9 ms and the levels of both target and preceding clicks were constant across contexts. Given this timing, contexts could differ 18 ms prior to target clicks. In order to examine the time window over which sound-level context affect human brainstem responses, we conducted Experiments III (N=16) and IV (N=19) using the same experimental design as in Experiment II (Figure 2A-C), but with onset-to-onset intervals of 14 ms and 22 ms, respectively.

Wave V latencies for the 25-dB and the 55-dB contexts for each of the three onset-to-onset intervals (9, 14, 22 ms) are shown in Figure 3A. Mean latency, independent of context, decreased with increasing onset-to-onset interval (9 ms vs. 14 ms: t_32_ = 4.392, p = 1.1×10^-4^, r_e_ = 0.613; 9 ms vs. 22 ms: t_35_ = 6.266, p = 3.4×10^-7^, r_e_ = 0.727; Figure 3B), indicating more adaptation of brainstem responses for short intervals between clicks (see also Experiment I and Supplementary Experiments A and B^28,31,41^).

**Figure 3:**
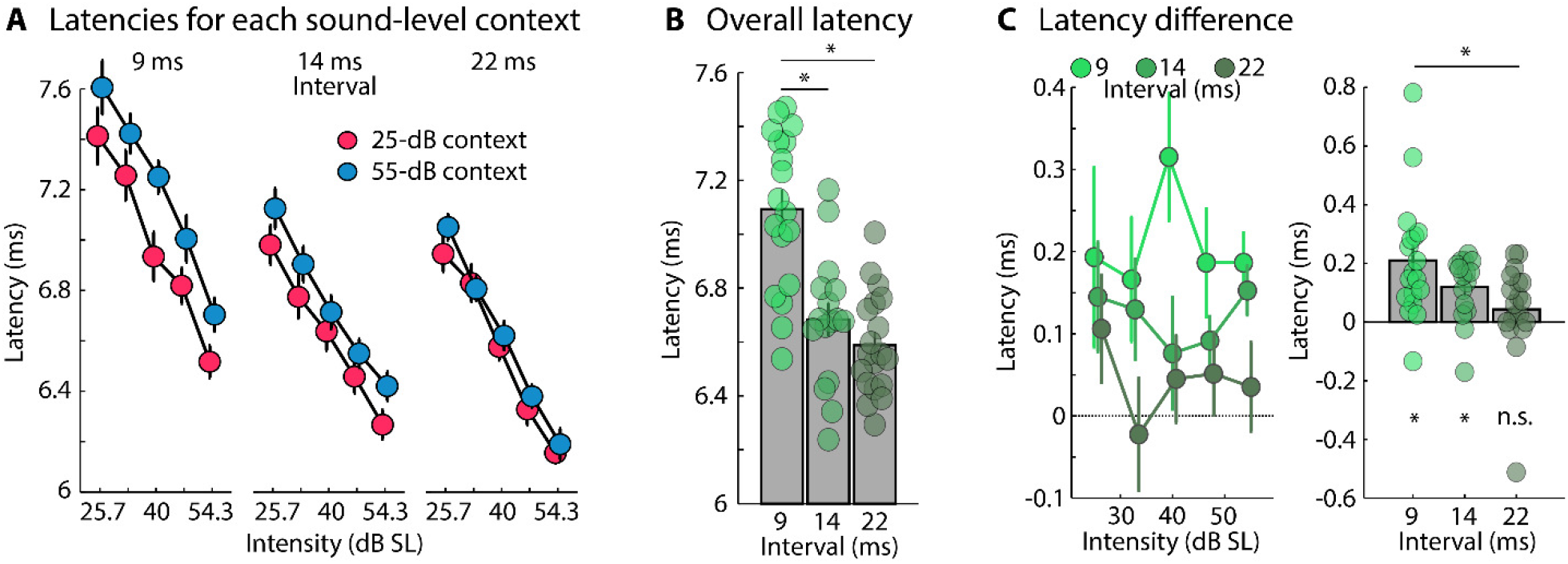
Neural adaptation to sound-level context for different click presentation rates. **A:** Wave V latencies for target clicks with different sound levels in two sound-level contexts (25-dB SL and 55-dB SL contexts) and for different click presentation rates (9 ms [Experiment II], 14 ms [Experiment III], 22 ms [Experiment IV]). **B:** Overall Wave V latency for target clicks presented at different presentation rates, independent of sound level and context. **C:** Latency difference between sound-level contexts (55-dB context minus 25-dB context) for each target click sound level (left) and averaged across target click sound levels (right). Asterisks below the bar graphs indicate a significant difference from zero (i.e., between the two contexts). Error bars reflect the standard error of the mean. *p ≤ 0.05, n.s. – not significant.

The difference between sound-level contexts (55-dB minus 25-dB context) was significant for 9 ms (t_17_ = 4.252, p = 5.4×10^-4^, r_e_ = 0.718) and 14 ms (t_15_ = 4.381, p = 5.4×10^-4^, r_e_ = 0.749) onset-to-onset intervals, but not for 22 ms onset-to-onset interval (t_18_ = 1.163, p = 0.260, r_e_ = 0.264; Figure 3C); although the absence of a context effect for the 22-ms interval appears to be driven by one outlier data point (Figure 3C, right), as its removal (t_17_ = 3.428, p = 0.003, r_e_ = 0.639) and a non-parametric test (p = 0.049, df = 18) revealed a significant difference. The effect of sound-level context (55-dB minus 25-dB context) was larger for the 9 ms compared to the 22 ms onset-to-onset interval (t_35_ = 2.721, p = 0.01, r_e_ = 0.418; Figure 3C; also when the outlier was removed from the 22-ms condition: t_34_ = 2.521, p = 0.017, r_e_ = 0.397), but did not differ between onset-to-onset intervals of 9 ms and 14 ms (t_32_ = 1.554, p = 0.13, r_e_ = 0.265; Figure 3C) or 14 ms and 22 ms (t_33_ = 1.601, p = 0.119, r_e_ = 0.269; Figure 3C).

The results from Experiments II-IV demonstrate that human brainstem responses are sensitive to sound-level context, but also that this sensitivity decreases when sounds are more spread out in time (i.e., a smaller context effect for the 22-ms than the 9-ms onset-to-onset interval condition). That brainstem responses were sensitive to sound-level context for the 22-ms interval (when the outlier was removed) – that is, to contextual differences in sound levels 44 ms or further in the past – suggests that sound-level information over at least a period of 44 ms are represented in human brainstem responses. This is also consistent with the identified temporal evolution of neural adaptation observed in Experiments I and Supplementary Experiments A and B (i.e., integration of approximately 40 ms of stimulation history).

## Discussion

In the current study, we conducted a series of six electroencephalography experiments (two are presented in the supplementary materials) to investigate the extent to which neural units in the human brainstem represent the history of acoustic stimuli, whether the human brainstem is sensitive to sound-level context of acoustic environments, and the timescale over which sound-level information is integrated by the brainstem. We show that acoustic stimuli within a time window of several tens of milliseconds are represented in human brainstem responses through neural adaptation (Experiment I and Supplementary Experiment A), while we exclude prediction-related processes as an underlying mechanism (Supplementary Experiment B). We further demonstrate that brainstem circuits represent sound-level context (Experiment II-IV) and that the sensitivity to sound-level context decreases as stimuli are more spread out in time. Our results provide information about the timecourses of neural adaptation in the human brainstem and are an important link to studies on neural adaptation in non-human animals.

### Human brainstem responses are sensitive to stimulation history spanning several sounds

Adaptation to sound-level context or other properties of the acoustic environment – often referred to as adaptation to stimulus statistics^15,16,18^ – requires that neurons represent stimulation history across several stimuli. For Experiment I and Supplementary Experiments A and B, we adapted experimental paradigms from our previous work on longer-term adaptation in the human auditory cortex^42,43^ to investigate the extent to which stimulation history is reflected in adaptation of human brainstem responses. We observed that the Wave V of the auditory evoked potential – originating from the brainstem^41,44–47^ – peaked later when more clicks occurred within a preceding time window (“decelerating” sequence) compared to fewer clicks (“accelerating” sequence), despite an identical interval directly preceding each target click and identical sound levels (Experiments I and Supplementary Experiment A; Figures 1 and S1). We estimate that, given the current stimulation protocols, acoustic stimuli over a period of several tens of milliseconds (^~^40 ms) are represented in human brainstem responses (e.g., Figure 1F). Finally, showing that the longer-term stimulation history affects the Wave V latency in sequences with clicks occurring at random, unpredictable times (Supplementary Experiment B), rules out that prediction-related processes underlie the changes in Wave V latency. Our data are more consistent with neural adaptation by, for example, affecting the spike latency of brainstem neurons^6,52^.

Our results demonstrating that Wave V latency is sensitive to stimulation history are consistent with previous work, using isochronous click sequences and click trains, showing that Wave V latency increases with increasing click rate^38,41^ and that Wave V latency is affected by the stimulation history spanning several sounds^28,35,39^. The degree to which the longer-term stimulation history modulates Wave V latency varies across studies^28,35,36,40^ and appears to depend to some extent on the sound level at which acoustic stimuli are presented^35,41^. This previous work also indicates that louder sounds have a longer-term influence on the latency of human brainstem responses than softer sounds^35^. Given this prior work, we expected to observe adaptation that reflected stimulation history in the current study. However, the variability in previous work and the potential dependence on experimental parameters, such as sound level, required a detailed investigation in Experiment I and Supplementary Experiments A and B to establish effects of stimulation history before investigating adaptation to sound-level context. We show here that brainstem responses represent the acoustic stimulation history in accelerating-decelerating sequences and random-interval sequences for several tens of milliseconds.

### Human brainstem responses adapt to sound-level context

In three EEG experiments, we investigated the extent to which neurons in the human brainstem adapt to sound-level context. We utilized a paradigm in which participants listened to clicks in two contexts, where the sound level was drawn from distributions with different modal sound levels (25 dB SL and 55 dB SL) and where target clicks and clicks preceding target clicks were identical in the two contexts with different modal sound levels^16,24^. Controlling the immediate history of clicks across contexts allowed us to investigate the effect of longer-term sound-level context on brainstem responses. We observed that Wave V latency, but not amplitude was affected by sound-level context, such that Wave V peaked later in the 55-dB context compared to the 25-dB context (Figures 2 and 3). We show that the sensitivity of the human brainstem response to sound-level context decreases when the time window increases over which acoustic stimuli need to be integrated, and that this context effect was present when at least 44 ms of sound stimulation prior to target clicks was identical in both contexts (Figure 3C; Experiment III; 22-ms onset-to-onset interval).

Our observation of sensitivity to sound-level context in the human brainstem is consistent with previous work showing adaptation to sound-level context in rodent inferior colliculus^15,18,21,22^ and auditory nerve fibers^19,20^, and with studies investigating the human auditory cortex^16,24–27^. Our findings also complement a broader literature showing adaptation to other features and distributions of acoustic environments, in addition to sound level, such as acoustic contrast^53,54^, mean and variance of interaural level^55^, spectral variance^6,12,13^, and other complex distributional properties^14^. Critically, the current study ensured that the N-1 history is controlled across our two sound-level contexts. Most previous work investigating adaptation to contextual (or statistical) information did not control for the immediate stimulation history^15,18–21^, and so in these studies, effects of context may in part be driven by the local stimulation history.

The term adaptation to stimulus statistics or sound-level statistics is commonly used to refer to altered responses with experience to auditory stimuli in different stimulation contexts as here^15,18–20,56^. The term may imply that neuronal populations code for probabilities of acoustic features and modulate their responses according to the median, mean, or other properties of acoustic dimensions^14,15,55^. However, several putative mechanisms, such as synaptic depression and inhibition, can lead to adaptation^5,57^ without necessitating estimations of properties of acoustic dimensions – and thus stimulus statistics – in a strict sense. The inter-stimulus intervals utilized here, however, may suggest that refractoriness of neurons contributes less to the current results since refractory periods are typically short (<5 ms). We show in Experiments 1 and Supplementary Experiments A and B that human brainstem responses are sensitive to the history of interstimulus intervals, and that the adaptation timescale in these experiments appears similar to the timescale we observed for sound-level adaptation in Experiments II-IV. This result may suggest that the same mechanism, for example, reflecting adaptation to sound energy levels integrated over time, may have been recruited during stimulation in all six experiments of the current study.

Previous work suggests that timescales of adaptation increase along the auditory pathway, with shorter timescales in the brainstem compared to the auditory cortex^54^. This is consistent with our observation of a reduction in sensitivity of human brainstem responses to sound-level context over a few tens of milliseconds (Figure 3), and previous work that demonstrates that the human auditory cortex shows adaptation lasting for 10 s or more^9,10,30^.

One difference between the current work and previous work is that we show adaptation of response latency, but not amplitude, to sound-level context (Figure 2H). For example, Wave V amplitude was not sensitive to sound-level context (although it was sensitive to sound level itself). In contrast, previous investigations of adaptation to sound-level statistics report sensitivity of firing rate in non-human mammals or magneto-/electroencephalographic response magnitude in humans^15,16,18,21,22,24^, whereas no effects of spike latency or response latency were reported; but see^6^. In these previous studies, sensitivity of firing rate and field potential magnitude appeared to shift such that differences in responses between sound-level contexts emerged most strongly for sound levels around the modal level, whereas responses were absent for low-level sounds and saturated for high-level sounds^15,16,18–22^. The current study, in contrast, shows an overall shift in response latency for all levels tested. Whether our effect of sound-level context on response latency is consistent with the work in animals is unknown, because latency has not been investigated previously.

Although it is well known that Wave V amplitude adapts less than Wave V latency^32,36^, the reasons for this are unknown. One previous study revealed frequency-specific adaptation of spike latency in the inferior colliculus of rats, but no difference in latency adaptation between two contexts differing in spectral variance^6^. However, in this previous study, stimulus presentation was much slower (at an onset-to-onset interval of 300 ms) and so no context effect on latency would be expected, based on the current results. Future work in rodents may investigate more closely whether spike latency also adapts to sound-level context to enable more direct comparisons with the current study.

## Conclusions

In the current study, we investigated the extent to which neural responses in the human brainstem adapt to the acoustic stimulation history beyond the preceding stimulus and whether the sound-level context is represented in the response. We also explored over which timescale sound-level information is reflected in human brainstem activity. We demonstrate that acoustic stimuli over several tens of milliseconds (^~^40 ms) are represented in the latency of human brainstem responses, likely via neural adaptation mechanisms but not through prediction-related processes. We further show that brainstem responses represent sound-level information up to at least 44 ms, and that this neural sensitivity to sound-level context increases with increasing density of the acoustic stimulation. Adaptation to sound-level context may support sound-level perception in dynamically changing acoustic environments. Our results demonstrate the timescale of neural adaptation in the human brainstem and provide a link to studies on neural adaptation in non-human animals.

## Methods and Materials

### Consent and ethics approval

A hundred and twenty-eight normal hearing adults (17–32 years) participated in the six electroencephalography experiments of the current study. Demographic information about participants is provided separately for each experiment. Four of the experiments are detailed here, whereas two others – Supplementary Experiments A and B – are presented in the supplementary materials. Participants gave prior written informed consent and were paid $5 CAD per half-hour for their participation or received course credits. The study was conducted in accordance with the Declaration of Helsinki, the Canadian Tri-Council Policy Statement on Ethical Conduct for Research Involving Humans (TCPS2-2014), and was approved by the Nonmedical Research Ethics Board of the University of Western Ontario (protocol ID: 106570). Participants reported no neurological disease or hearing problems and were naïve to the purposes of the experiment.

### General procedure and acoustic stimulation setup

Sounds used in the current series of experiments consisted of clicks of 0.1 ms duration (rectangle function) presented at different onset-to-onset intervals and sound levels depending on the specific experiment described below. Click polarity was inverted on half of the click presentations and polarity inversion was tightly controlled across conditions within each experiment. Clicks were presented to the right ear via an Etymotic ER1 insert earphone, using a Fireface 400 external sound card controlled by Psychtoolbox (version 3.14) in MATLAB (MathWorks Inc.). The sensation level for click trains was determined for each participant using a method of limits procedure^58^, and all experimental stimuli were presented relative to this sensation level (see details below).

### Electroencephalographic recordings and preprocessing

Electroencephalographic activity was recorded using a 16-channel BioSemi (Active 2) system. Additional electrodes were placed at the left and right mastoids. EEG was recorded at 16,384 Hz with an online low-pass filter of 3,334 Hz. Electrodes were referenced online to a monopolar reference feedback loop connecting a driven passive sensor and a common-mode-sense (CMS) active sensor, both located posteriorly on the scalp.

MATLAB software (v7.14; MathWorks, Inc.) was used for offline data analysis. Raw data were filtered with an elliptic filter to suppress line noise at 60 Hz. Data were re-referenced by averaging the two mastoid channels and subtracting the average separately from each of the 16 channels. Data were high-pass filtered at 130 Hz (2743 points, Hann window) and low-pass filtered at 2500 Hz (101 points, Hann window). Data were divided into epochs ranging from 2 ms pre-click-onset to 10 ms post-click onset. Epochs were excluded if the signal change exceeded 60 μV in the 0 to 10 ms time window in either the Cz or C3 electrode. These electrodes were of particular interest because they are known to capture the Wave V most strongly, given right ear acoustic stimulation^59^.

The peak latency of the Wave V for each condition was identified as the latency at which the amplitude was highest within a 1.6 ms window centered on the average Wave V latency across conditions. We confirmed through visual inspection that latencies were correctly identified. The same automated and visual confirmation procedures were used for all experiments. Wave V amplitude was calculated by averaging the amplitude within a 0.2 ms time window centered on Wave V latency.

### Experiment I

#### Participants

Seventeen individuals participated in Experiment I (age range: 18–28 years; median age: 22 years; 11 females, 6 males). Data from three additional participants were recorded but excluded at the analysis stage, because Wave V was absent (N=2) or an atypically late Wave V latency was exhibited, peaking later than 8.5 ms (N=1). Results were unaffected by inclusion vs. exclusion of the latter participant.

#### Acoustic stimulation

Participants listened to six blocks of stimulation (each about 8 min long) that comprised click-train stimuli, in which the onset-to-onset interval of clicks changed from 60 ms to 10 ms (in 7 logarithmically-spaced steps) and back from 10 ms to 60 ms (in 7 logarithmically-spaced steps), creating an accelerating-decelerating stimulation cycle (Figure 1A; see also^42,43^). Across the six blocks, participants listened to 7,200 accelerating-decelerating stimulation cycles. All clicks were presented at 60 dB SL.

#### Data analysis

Single trials were separated into an ‘accelerating’ context (60 ms to 10 ms) and a ‘decelerating’ context (10 ms to 60 ms). As such, each context (accelerating, decelerating) comprised the same onset-to-onset intervals and, consequently, any response differences between contexts cannot be attributed to the duration of the interval directly preceding a click, but must be attributed to the longer-term stimulation history (in which the decelerating context compared to the accelerating context comprised more clicks in the same preceding time). Single-trial time courses for a unique onset-to-onset interval and one of its direct neighbors (i.e., shorter and longer intervals) were binned and averaged^43,60^. The Wave V latency and amplitudes were calculated for each onset-to-onset interval and each context (accelerating, decelerating). For statistical analyses, repeated-measured analyses of variance (rmANOVA) with the factors Interval and Context (accelerating, decelerating) were calculated separately for the Wave V latency and the Wave V amplitude.

### Experiment II

#### Participants

Eighteen individuals participated in Experiment II (age range: 17–32 years; median age: 18 years; 15 females, 3 males). None of them had participated in Experiment I. Data from two additional participants were excluded because of large stimulus artifacts in the recordings (N=1) and the absence of an identifiable Wave V (N=1).

#### Acoustic stimulation

The experimental paradigm was derived from our previous work on neural adaptation to sound-level context in the auditory cortex^16,24^. Clicks were presented in two types of contexts that differed with respect to the sound-level distributions from which a click’s sound level was drawn: for one context, the modal sound level was 25 dB SL, for the other context it was 55 dB SL (Figure 2A). To investigate the effects of sound-level context on human brainstem responses, we eliminated the confounding effects of different acoustics by ensuring that the clicks for which we analyzed neural responses as well as their local context (i.e., directly preceding clicks) were identical across contexts (i.e., 25 dB SL vs. 55 dB SL), whereas the longer-term sound-level information were different between the two contexts.

Participants listened to six blocks, each approximately 7 min long. In each block, 44,800 clicks were presented at a constant onset-to-onset interval of 9 ms. In each block, eight target clicks with sound levels ranging from 15 dB SL to 65 dB SL (step size: 7.143 dB SL) were each presented 960 times, yielding 7,680 target clicks per block. These stimuli were identical across all six blocks (black dots in Figure 2B/C). The 7,680 clicks immediately preceding each of these 8 · 960 = 7,680 target clicks were also fixed, such that for each of the 960 presentations of one of the eight sound levels, the sound level of the preceding tone took on one of 30 sound levels (range: 15 dB SL to 65 dB SL; step size: 1.724 dB SL) without replacement. Thus, the same 7,680 pairs of experimental clicks were presented in each block.

The sound levels for the remaining 29,440 clicks (out of the 44,800 clicks per block) were chosen randomly (range: 15 dB SL to 65 dB SL; step size: 0.1 dB SL) depending on the sound-level distribution (Figure 2A). For three out of the six blocks, sound levels were randomly chosen from a sound-level distribution with a 25-dB high-probability region (Figure 2B). For the other three blocks, sound levels were randomly chosen from a sound-level distribution with a 55-dB high-probability region (Figure 2C). High probability regions had a width of 10 dB centered on 25 dB SL or 55 dB SL (Figure 2A-C). The 15,360 experimental clicks (i.e., 7,680 pairs) and the 29,440 filler clicks were randomly intermixed in each block and presented such that at least one filler click occurred between each of the 7,680 pairs of experimental clicks. Blocks with different sound-level contexts alternated, and the starting sound-level context was counter-balanced across participants. Analysis of the influence of sound-level context on neural responses focused on the second click in each pair of experimental clicks: these were identical across the two sound-level distributions (black dots in Figure 2B/C).

#### Data analysis

The number of trials for each of the 8 sound levels in the different contexts was relatively low for the analysis of the small Wave V brainstem potential (2880 trials = 960 trials × 3 blocks). In order to increase the number of trials in the response average, thereby increasing the signal-to-noise ratio^16,60^, we averaged trials for four adjacent sound levels out of the 8 sound levels (separately for the 25-dB SL context and the 55-dB SL context). For example, single trials for sounds with the 15, 22.1, 29.3, and 36.4 dB SL sound level were averaged. Averaging trials for four adjacent sound levels reduced the number of unique sound levels from 8 to 5, while increasing the number of trials in a unique response average. Our choice of smoothing across different sound-level conditions also ensured that a sufficiently high number of trials were averaged in Experiments III and IV, in which longer onset-to-onset intervals reduced the number trials per sound level that were presented (see below). Wave V latency was extracted for each sound level and context, and a rmANOVA with factor Sound Level and Context (25-dB SL, 55-dB SL) was calculated.

### Experiment III

#### Participants

Sixteen individuals participated in Experiment III (age range: 18–32 years; median age: 18.5 years; 12 females, 4 males). None of them had participated in Experiment I or II.

#### Acoustic stimulation

Stimulation procedures were identical to Experiment II with the following exceptions. Participants listened to six blocks, each approx. 8.5 min long. In each block, 36,400 clicks were presented at a constant onset-to-onset interval of 14 ms. 6,240 experimental click pairs (8 sound levels × 780 trials) and 23,920 filler clicks were presented per block.

#### Data analysis

The same analyses as for Experiment II were carried out.

### Experiment IV

#### Participants

Nineteen individuals participated in Experiment IV (age range: 17–23 years; median age: 18 years; 13 females, 5 males; one person did not provide information about their sex). None of them had participated in any of the other experiments.

#### Acoustic stimulation

Stimulation procedures were identical to Experiment II with the following exceptions. Participants listened to six blocks, each approx. 8.3 min long. In each block, 22,400 clicks were presented at a constant onset-to-onset interval of 22 ms. 3,840 experimental click pairs (8 sound levels × 480 trials) and 14,720 filler clicks were presented per block.

#### Data analysis

The same analyses as for Experiment II were carried out.

### Effect sizes

Throughout the manuscript, effect sizes are provided as partial eta squared (η_p_^2^) for repeated-measures ANOVAs and as r_e_ (r_equivalent_) for t-tests^61^.

## Acknowledgements

BH was supported by a BrainsCAN postdoctoral fellowship (Canada First Research Excellence Fund; CFREF) and by the Canada Research Chair program.

## Author contributions

The experiments were performed in the Brain and Mind Institute at the University of Western Ontario, London, Canada. B.H., S.Y.., I.S.J. designed the study. B.H., S.J., K.A. recorded the data. B.H. programmed the experiments and analyzed the data. S.Y. and D.P. supported data analysis. B.H., S.Y., D.P., and I.S.J. interpreted the results. B.H. wrote the manuscript. All authors reviewed and revised the manuscript.

## Data availability

The datasets generated during the current study are not publicly available because participant consent for public-data sharing was not obtained. Data are available from the corresponding author on reasonable request.

