## Supplemental Materials for "Sound level context modulates neural activity in the human brainstem"

### Supplementary Results

#### *Human brainstem activity is affected by the longer-term history of acoustic stimulation*

In Supplementary Experiment A, normal-hearing younger adults (N=28; data reanalyzed from Yasmin et al., 2020) listened to click sequences presented at 60 dB sensation level (SL) that continuously accelerated and decelerated by changing the click onset-to-onset interval from 40 ms to 4 ms and back to 40 ms (spaced logarithmically; Figure S1A). Analyses focused on the Wave V potential (Figure S1B), which is known to originate from the brainstem (Don et al., 1977; Møller and Jannetta, 1982; Møller and Jannetta, 1983; Møller, 1998; Parkkonen et al., 2009). Wave V amplitude and latency were calculated, separately for clicks preceded by different onset-to-onset intervals. Wave V latency decreased with increasing duration to a previous click ($t_{27}$ = -6.430, p = 6.9×$10^{-7}$, $r_e$ = 0.778; Figure S1C, right), whereas Wave V amplitude was not affected ($t_{27}$ = 0.194, p = 0.848, $r_e$ = 0.037; Figure S1C, left). This analysis demonstrates that the immediate stimulation history affects the latency of the brainstem response.

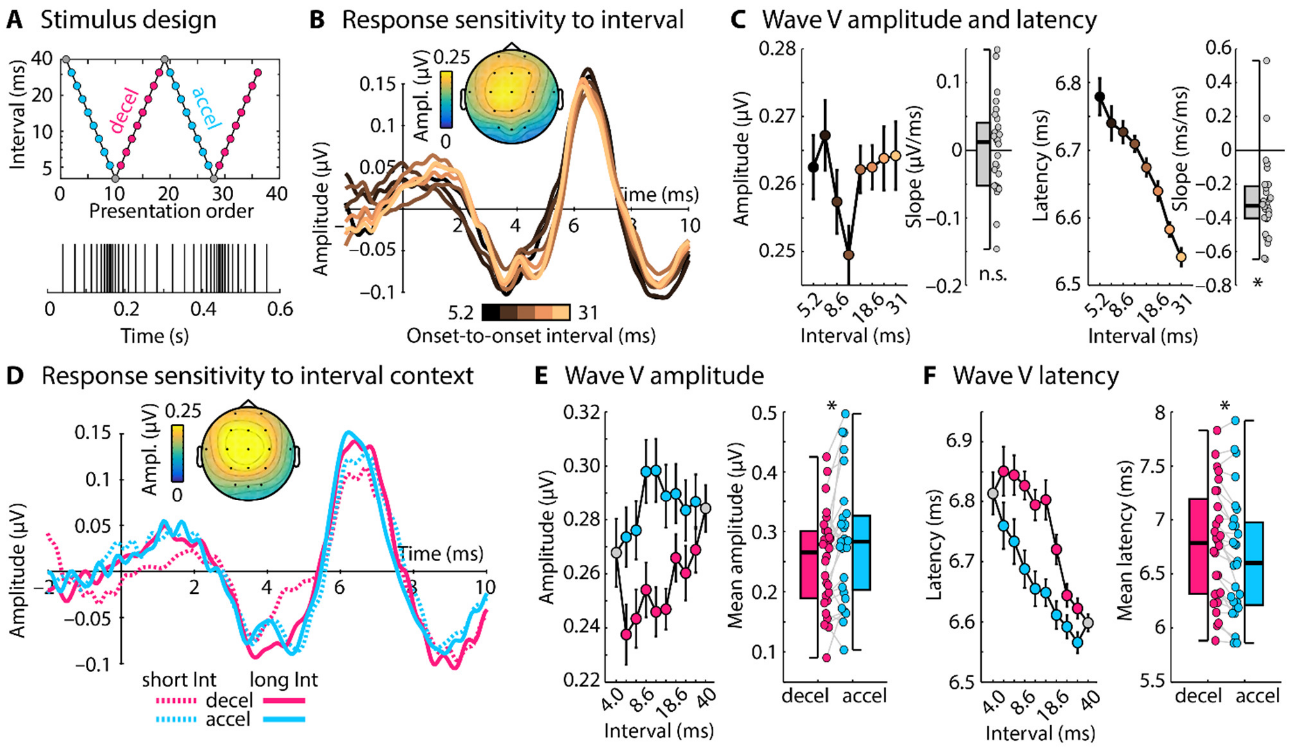


**Figure S1: Stimulus design and results for Supplementary Experiment A. A:** Onset-to-onset intervals (logarithmic y-axis) of accelerating and decelerating click sequences. **B:** Response time courses to clicks, separately for different onset-to-onset intervals to a preceding click. The topographical distribution reflects the mean Wave V (~6.6 ms). **C:** Wave V amplitude and latency as a function of onset-to-onset interval. Box plots and individual data points (dots) reflect the slope from a linear function fit relating Wave V amplitude/latency to onset-to-onset intervals. Wave V latency, but not amplitude, was sensitive to the inter-click-interval (slope significantly different from zero, $p \le 0.05$). **D:** Response time courses for a short (6.7 ms) and a long interval (24 ms), separately for decelerating and accelerating sequence contexts. Topographical distribution reflects the mean Wave V. **E/F:** Wave V amplitude and latency as a function of onset-to-onset interval, separately for decelerating and accelerating contexts. Box plots and individual data points (dots) reflect the mean difference between amplitudes/latencies for decelerating and accelerating contexts. Amplitudes were larger and latencies smaller in accelerating compared to decelerating contexts ($p \le 0.05$), demonstrating longer-term influences of the acoustic stimulation on the Wave V response. Error bars reflect the standard error of the mean (removal of between-subject variance; Masson and Loftus, 2003). *$p \le 0.05$, n.s. – not significant

In order to investigate the influence of the longer-term stimulation history – beyond the interval directly preceding a click – on brainstem responses, trials were separated into accelerating and decelerating sequence contexts (Figure S1D). Both accelerating and decelerating contexts comprise the same onset-to-onset intervals and, as a result, any differences in brainstem responses between contexts could only arise from the longer-term stimulation history: clicks in the decelerating context are preceded by more clicks in same time window compared to the accelerating context, despite the same interval duration directly preceding a click. Wave V amplitude (averaged across onset-to-onset intervals) was larger ($t_{27} = 2.770$, $p = 0.010$, $r_e = 0.471$; Figure S1E) and Wave V latency smaller in the accelerating compared to the decelerating context ($t_{27} = 3.949$, $p = 5.1\times10^{-4}$, $r_e = 0.605$; Figure S1F). Previous work suggests that the morphology of the auditory brainstem response can be affected by overlapping responses for shorter intervals (Scott and Harkins, 1978). Waveforms in Figure S1B provide no indication of large morphological changes. This could potentially be due to our non-isochronous stimulus paradigm compared to the isochronous stimulation in previous work (Scott and Harkins, 1978). Critically, we also observed effects of context when intervals shorter than 8.5 ms, which could potentially lead to responses interfering with the Wave V at ~6.6 ms, were excluded from the analysis (amplitude: $t_{27} = 2.137$, $p = 0.042$, $r_e = 0.380$; latency: $t_{27} = 3.560$, $p = 0.001$, $r_e = 0.565$).

***Human brainstem activity is affected by longer-term stimulation history in click sequences with unpredictable intervals***

Because click sequences in Experiment I and Supplementary Experiment A accelerated and decelerated in a regular, and thus predictable, fashion, it is unclear whether a prediction-related process or neural adaptation underlies the observed effects of stimulation history. If neural adaptation underlies the modulation of Wave V latency, but not a prediction-related process, we should also observe effects of the longer-term stimulation history on Wave V latency for clicks presented in sequences with random, unpredictable onset-to-onset intervals.

In Supplementary Experiment B, normal-hearing younger adults (N=19) were presented with sequences of 60-dB SL clicks in which the onset-to-onset interval between clicks varied pseudo-randomly between 4 ms and 40 ms (spaced logarithmically; Figure S2A). Wave V latency ($t_{18}$ = -3.775, p = 0.001, $r_e$ = 0.665), but not amplitude ($t_{18}$ = -2.074, p = 0.053, $r_e$ = 0.439), was sensitive to the duration of the interval directly preceding a click (Figure S2B,C).

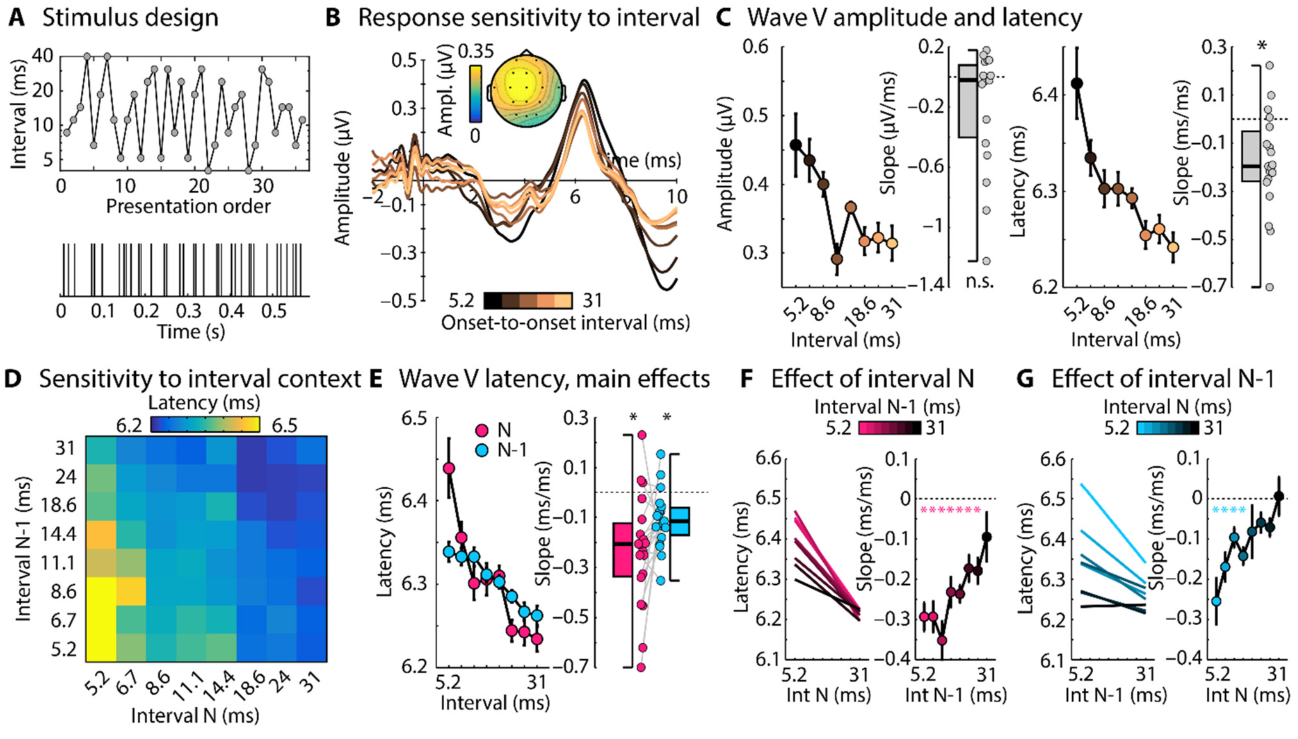


**Figure S2: Stimulus design and results for Supplementary Experiment B. A:** Stimulation comprised a click sequence in which the onset-to-onset interval of clicks varied randomly between 4 ms and 40 ms. **B:** Response time courses to clicks were binned according to the time interval directly preceding a click. The topographical distribution reflects the mean Wave V. **C:** Wave V amplitude and latency as a function of inter-click-interval. Box plots and individual data points (dots) reflect the slope from a linear function fit relating Wave V amplitude/latency to onset-to-onset intervals (separately for each participant). Wave V latency, but not amplitude, was sensitive to the inter-click-interval ($p \leq 0.05$). **D:** Wave V latency as a function of preceding interval N and N-1. **E:** Marginals of Wave V latency in Panel D. Box plots and individual data points (dots) reflect the slope from a linear function fit relating Wave V latency to onset-to-onset intervals, separately for intervals N and N-1. Latencies decreased with increasing interval duration for both N and N-1 intervals ($p \leq 0.05$). **F:** Left plot shows the predicted values from linear function fits relating Wave V latency to intervals N, separately for all penultimate intervals N-1. Right plot shows the respective mean slopes from the fits. An asterisk marks a significant difference from zero (FDR-thresholded). **G:** Left plot shows the predicted values from linear function fits relating Wave V latency to penultimate intervals N-1, separately for all intervals N. Right plot shows the respective mean slopes from the fits. An asterisk marks a significant difference from zero (FDR-thresholded). Error bars reflect the standard error (removal of between-subject variance; Masson and Loftus, 2003). *$p \leq 0.05$, n.s. – not significant

In order to analyze whether the longer-term stimulation history influences the Wave V latency to clicks in sequences with unpredictable intervals, we sorted each trial into a 2-dimensional grid according to the onset-to-onset interval directly preceding the click (interval N) and the penultimate interval (N-1). Trials were averaged and the Wave V latency was estimated for each grid point (Figure S2D). Latencies were longest when both interval N and N-1 were short (bottom left corner of Figure S2D), and shortest when both interval N and N-1 were long (top right corner of Figure S2D).

We analyzed the effects of the directly preceding interval (N) and the penultimate interval (N-1) by averaging Wave V latencies across the vertical and horizontal dimensions of the 2D grid in Figure S2D, respectively. A linear function was fit to these averaged Wave V latencies as a function interval duration, separately for N and N-1 intervals (Figure S2E). The mean slopes relating Wave V latency to intervals N ($t_{18} = -4.478$, $p = 2.9\times10^{-4}$, $r_e = 0.726$) and intervals N-1 ($t_{18} = -4.129$, $p = 6.3\times10^{-4}$, $r_e = 0.697$) were smaller than zero, demonstrating that it is not only the time interval of the directly preceding click that affects Wave V latency, but also the penultimate interval. Slopes did not differ between interval N and interval N-1, but trended towards a shallower slope for N-1 intervals ($t_{18} = 2.024$, $p = 0.058$, $r_e = 0.431$; Figure S2E). The pattern of results was also observed when trials whose subsequent interval was 8.5 ms or smaller, and that thus may lead to superimposed responses, were excluded from the analysis (slope for interval N: $t_{18} = -4.037$, $p = 7.7\times10^{-4}$, $r_e = 0.689$; slope for interval N-1: $t_{18} = -2.306$, $p = 0.033$, $r_e = 0.478$).

Figure S2F and S2G provide a more detailed depiction of the influence of sound history on brainstem responses. Figure S2F shows that the directly preceding interval (N) only influenced the Wave

V latency when the penultimate interval (N-1) was shorter than ~30 ms. This indicates that longer-term stimulation history beyond interval N is, in fact, needed to modulate Wave V latency. Figure S2G additionally shows that the penultimate interval (N-1) modulated the Wave V latency only when the interval N was 11 ms or shorter, but not when the interval N was 14 ms or above (FDR-thresholded).

The analyses of Experiment I and Supplementary Experiments A and B show that the latency of brainstem activity (Wave V) in response to a click is strongly influenced not only by the interval directly preceding a click, but also by clicks further in the past (see also Lasky et al., 1996; Lasky, 1997). Neural activity patterns of the human brainstem appear to integrate preceding sound information over at least 40 ms, given our recording and stimulation protocols. The irregular, unpredictable click stimulation used in Supplementary Experiment B excludes the possibility that the influence of stimulus history on brainstem responses is due to prediction-related processes. The data are more consistent with neural adaptation underlying changes in brainstem response latency.

### Supplementary Methods

#### *Supplementary Experiment A*

**Participants.** Twenty-eight individuals participated in Supplementary Experiment A (age range: 17–28 years; median age: 18 years; 20 females, 8 males). Data analyzed for this experiment have been published previously (Yasmin et al., 2020) and are re-analyzed here in a new way. None of the participants participated in any of the other experiments. Data from six additional participants were recorded but excluded at the analysis stage. Wave V was absent for two participants. Four participants exhibited an atypically late Wave V latency, peaking later than 8.5 ms. Results were unaffected by excluding the latter four participants.

**Acoustic stimulation.** Participants listened to six blocks of stimulation (each about 6 min long) that comprised click-train stimuli, in which the onset-to-onset interval of clicks changed from 40 ms to 4 ms and back to 40 ms, creating an accelerating-decelerating stimulation cycle (18 logarithmically spaced intervals; Figure S1A). Across the six blocks, participants listened to 2,304 accelerating-decelerating stimulation cycles. All clicks were presented at 60 dB SL. For additional details see Yasmin et al., 2020 (see also Herrmann et al., 2016; Herrmann et al., 2019).

**Data analysis.** Single trials were separated into an 'accelerating' context (40 ms to 4 ms) and a 'decelerating' context (4 ms to 40 ms). As such, each context (accelerating, decelerating) comprised the same onset-to-onset intervals and, consequently, any response differences between contexts cannot be attributed to the duration of interval directly preceding a click, but must be attributed to the longer-term stimulation history (in which the decelerating context compared to the accelerating context comprised more clicks in the same preceding time). Single-trial time courses for a unique onset-to-onset interval and two of its direct neighbors (i.e., shorter and longer intervals) were binned and averaged (Ingham and McAlpine, 2005; Herrmann et al., 2016). The Wave V latency and amplitudes were calculated for each onset-to-onset interval and each context (accelerating, decelerating). For statistical analyses, the mean Wave V latency and amplitude across onset-to-onset intervals was calculated separately for clicks presented in the accelerating and decelerating contexts (excluding the 4 ms and 40 ms intervals). Differences between contexts were assessed using dependent-samples t-tests.

*Supplementary Experiment B*

**Participants.** Nineteen individuals participated in Supplementary Experiment B (age range: 17–23 years; median age: 18 years; 11 females, 8 males). None of them participated in any of the other experiments.

**Acoustic stimulation.** Participants listened to six blocks of stimulation (each about 7 min long) in which click stimuli were presented pseudo-randomly at 10 onset-to-onset intervals ranging from 4 ms and 40 ms (logarithmically spaced intervals; Figure S2A). 2500 clicks per block of stimulation were presented for each of the 10 onset-to-onset intervals, resulting in 12,500 clicks for each onset-to-onset interval over the whole experiment. All clicks were presented at 60 dB SL.

**Data analysis.** Single trials were grouped into a two-dimensional grid according to the directly preceding interval N and the penultimate interval N-1 of a click. In order to increase the signal-to-noise ratio, single-trial time courses for a unique interval and its direct neighbors (i.e., shorter and longer intervals) were grouped into one grid bin and averaged (Ingham and McAlpine, 2005; Herrmann et al., 2016). The Wave V latency was calculated for each N by N-1 interval grid point. We analyzed the effects of the directly preceding interval (N) and the penultimate interval (N-1) on Wave V latency by averaging latencies across the intervals N-1 and intervals N, respectively (i.e., vertical and horizontal dimensions, respectively, of the 2D grid in Figure S2D). For each participant, a linear function was fit to the averaged Wave V latencies as a function of interval duration, separately for N and N-1 intervals. The estimated linear coefficients resulting from the fit were tested against zero using a one-sample t-test, separately for N and N-1 intervals.

In order to obtain a more detailed picture of the influence of the directly preceding interval (N) and the penultimate interval (N-1) on Wave V latency, a linear function fit was used to relate Wave V latency to intervals N, separately for each interval N-1, and to relate Wave V latency to intervals N-1, separately for each interval N. The resulting linear coefficients were tested against zero using a one-sample t-test. False discovery rate (FDR) was used to correct for multiple comparisons (Benjamini and Hochberg, 1995; Genovese et al., 2002).
